# TMEM Doorway Mediated Metastasis in Pancreatic Ductal Adenocarcinoma by Tie2 Signaling

**DOI:** 10.64898/2025.12.05.692608

**Authors:** Erika Pereira Zambalde, Lina A. Ariyan, Francisco Puerta-Martinez, Yu Quin Zhu, Priyanka Patil, Nicole C. Panarelli, Jiufeng Li, Yookyung Jung, Christian Adkisson, Jakeb Petersen, Dianne Cox, Hava Gil-Henn, Robert Eddy, Ben Z. Stanger, David Entenberg, Maja H. Oktay, John S. Condeelis, John C. McAuliffe

## Abstract

Pancreatic ductal adenocarcinoma (PDAC) is almost invariably fatal due to early hematogenous dissemination that occurs before the primary tumor is clinically detectable, yet the cellular mechanism of tumor cell intravasation has remained unknown. Using multiphoton intravital imaging in autochthonous and orthotopic PDAC models, we demonstrate that intravasation occurs at Tumor Microenvironment of Metastasis (TMEM) doorways—tri-cellular structures comprising a MENA-expressing tumor cell, a Tie2⁺ macrophage, and an endothelial cell in direct contact. These structures are abundant in human PDAC, enriched for Tie2⁺ macrophages, and markedly reduced after neoadjuvant chemotherapy. Selective pharmacologic inhibition of Tie2 with rebastinib decreases TMEM-associated transient vascular openings, suppresses circulating and hepatic disseminated tumor cells, and—when combined with perioperative FOLFIRINOX after curative-intent resection—improves median survival in murine PDAC. These findings establish TMEM doorways as a common, druggable mechanism of intravasation across epithelial cancers and identify Tie2⁺ macrophages as a therapeutic target to prevent metastatic seeding in PDAC, a disease with no anti-metastatic therapies. TMEM doorway-mediated intravasation in PDAC supports its role as a common gateway for hematogenous metastasis in carcinoma.

## Introduction

Pancreatic ductal adenocarcinoma (PDAC) remains among the most lethal solid tumors, with a 5-year survival rate of less than 10%, despite ongoing advancements in diagnostic and therapeutic modalities (1). Most patients present with either locoregional or limited metastatic disease. However, despite maximal multimodality therapies, the disease progresses to overwhelming metastatic burden. Thus, the poor prognosis of PDAC is related to overwhelming metastatic burden. Therapeutics designed to impede the progression of PDAC from locoregional or limited metastatic burden to overwhelming metastasis are needed to improve the mortality rate of PDAC.

Metastasis is a multistep process including invasion, intravasation, survival in circulation, extravasation, and colonization of distant organs (2). Among these, intravasation of tumor cells into the vasculature is a crucial, yet poorly understood, event in PDAC metastasis. The intravasation event could be a random result of tumor cell migratory and invasive characteristics. Alternatively, the intravasation event could be a discrete tumor microenvironment signaling mechanism. Early histopathological analyses of resected PDAC specimens identified tumor cells near vascular spaces but lacked the temporal resolution to capture dynamic intravasation events (3, 4). In addition, recent studies suggested that intravasation is not solely mediated by tumor-intrinsic mechanisms but also relies on interactions within the tumor microenvironment (TME) (3, 4).The PDAC TME is characterized by an extensive fibrotic stroma and a rich immune infiltrate, prominently featuring tumor-associated macrophages (TAMs), which are recognized as key facilitators of tumor progression, angiogenesis, immune suppression and metastasis (5).

The Tumor Microenvironment of Metastasis (TMEM) doorway has been identified as the mechanism of intravasation in breast cancer models. The TMEM doorway is a tri-cellular structure defined by a MENA^High^ tumor cell, Tie2⁺ macrophage, and endothelial cell in direct and stable contact. Signaling at the TMEM doorway leads to localized, vascular openings (TMEM doorway-associated vascular openings, TAVO) that facilitate tumor cell intravasation (6). TMEM doorways have provided mechanistic insights into how TAMs cooperate with tumor and endothelial cells to mediate intravasation of tumor cells. Using high-resolution intravital imaging, TAVO was observed, in conjunction with tumor cell intravasation (7). Mechanistically, paracrine signaling at TMEM doorways leads to large vascular openings which allow tumor cells to enter the blood stream (8, 9). The density of TMEM doorways in primary breast tumors correlates strongly with metastatic recurrence and has been clinically validated as an independent prognostic biomarker in patients with ER+/HER2– breast cancer (10, 11). A key signaling step in TMEM doorway function is the Tie2-mediated production of VEGF in the TMEM doorway macrophage which induces TAVO. Inhibition of Tie2 has been shown to block TMEM doorway TAVO and tumor cell intravasation in breast tumor mouse models (12). These data show a link between TMEM doorways and metastatic progression.

The intravasation event has not been observed in PDAC. We recently developed a stabilized window for intravital imaging of the murine pancreas (13, 14). We observed significant cellular motility in the stabilized murine pancreas. We therefore expected to be able to observe cellular migration, invasion, and intravasation of PDAC using this model.

Here, we employ intravital imaging in genetically engineered mouse models of PDAC to observe the intravasation event. Our real-time imaging studies reveal transient vascular opening events that co-localize with perivascular Tie2-expressing macrophages congruent with TAVO. We show that TMEM doorways are present and operational in PDAC. Moreover, we show that blockade of Tie2 signaling attenuates both vascular opening and tumor cell dissemination. These findings implicate the Tie2+ TAM and the TMEM doorway as actionable targets to mitigate dissemination in PDAC.

## Results

### Intravital Imaging Reveals Macrophage-Mediated Vascular Opening and Tumor Cell Intravasation in PDAC

We first sought to directly observe the migration and intravasation dynamics in PDAC with the hope to identify actional targets of metastasis. To do this we utilized our stabilized window for imaging the pancreas (SWIP) and extended time-lapse intravital microscopy (IVM). Using the YFP labeled KPC transgenic mouse intravenously injected with high molecular weight labeled dextran (HMWD) to distinguish tumor vasculature, intravital imaging of palpable pancreatic primary tumors revealed transient and localized extravasation of dextran. This indicated large vascular opening events (Figure 1A). These intra tumoral vascular openings were transient, resolving spontaneously over 10–30 minutes.

**Figure 1.**
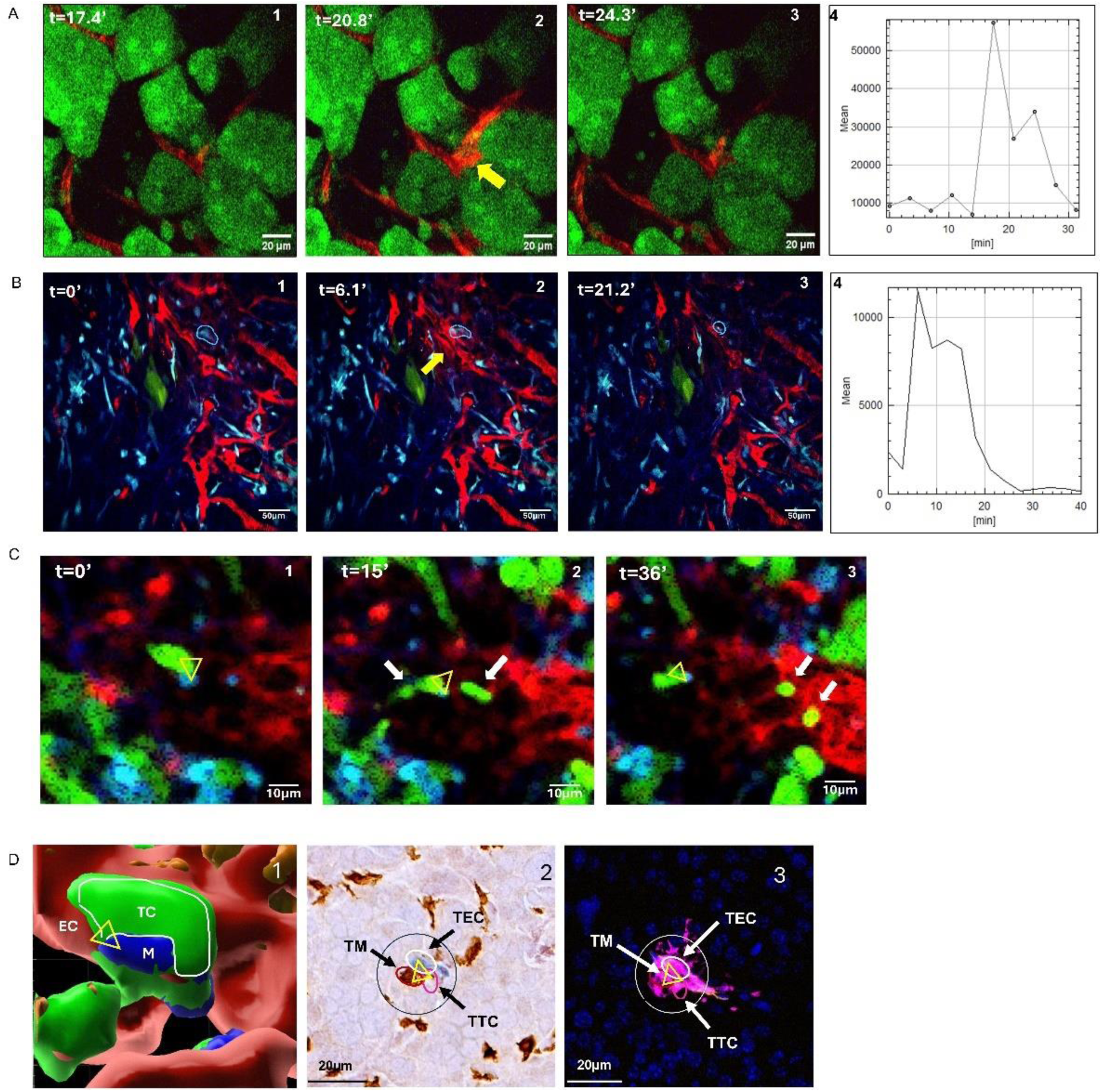
Intravital Imaging Reveals Macrophage-Associated Vascular Opening and Tumor Cell Intravasation in PDAC. (A) Time-lapse intravital microscopy in KPCY mice shows transient localized dextran extravasation (yellow arrow) in tumor regions (tumor cells: green).. (A1-3) panels showing timelapse images of the extravasation at the times indicated. (A4) quantifies the kinetics of dextran extravasation over time. (B) MACBlue mice imaged in an orthotopic YFP-labeled (green) PDAC tumor show dextran extravasation event (yellow arrow) associated with perivascular CFP+ macrophage (blue circle). (B1-3) panels showing timelapse images of the extravasation at the times indicated. (B4) Corresponding kinetics of dextran extravasation over time. (C) A series of frames captures a YFP+ tumor cell (green) interacting with a CFP+ macrophage (blue) near vasculature (red), followed by active intravasation into the bloodstream.. Panel 1 shows a TMEM doorway (triangle). Panel 2 shows two cancer cells intravasating (white arrows) at the TMEM doorway (yellow triangle). Panel 3 shows two tumor cells (whitearrows) that intravasated into bloodstream (white arrows) while the cells of the TMEM doorway remain in a stable tri-cell complex (yellow triangle). (D) Panel D1 shows a 3D intravital image of the tumor cell (TC, green), macrophage (M, blue) interaction when associated with a blood vessel endothelial cell (EC, red), identifying this stable three cell complex (triangle) as a TMEM doorway. D2 IHC stained formaldehyde fixed tissue section of the TMEM doorway (triangle). (D3) Image D2 overlaid with image of adjasent serial section showing extravasation of dextran occurs at the three cell complex confirming the three cell complex is a TMEM doorway.

Because we observed spatially distinct areas of HMWD extravasation within the tumor, we suspected this was related to the tumor microenvironment constituents other than tumor cells and endothelial cells. Previous reports suggest that TAMs are associated with metastasis (5, 15, 16). Using a syngeneic transgenic mouse model with CFP-labeled TAMs, we evaluated whether extravasation of HMWD was related to perivascular macrophages. We observed transient intra-tumoral opening at regions in the tumor vasculature with a macrophage positioned along the blood vessel (Figure 1B, yellow arrow). This observation suggested a macrophage-mediated tumor vascular opening mechanism.

We additionally observed YFP+ tumor cell streaming toward the tumor vasculature, engaging with a CFP+ perivascular macrophage at the vascular interface followed by intravasation of the tumor cells over approximately 30 minutes (Figure 1C). We only observed single cell migration and intravasation in tumors derived from KPCY C-EMT 3077 cells (Figure 1C). We did not observe collective migration or intravasating clusters in this model.

Three-dimensional reconstruction and modeling revealed a tri-cellular arrangement of macrophage, tumor cell, and endothelial cell at these intravasation sites (Figure 1D), consistent with a TMEM doorway (6). We therefore hypothesized that TMEM doorways are present and functional in PDAC.

### TMEM Doorways Are Abundant in Human PDAC and Associated with Aggressive Tumor Features

Based on our intravital imaging observations in PDAC mouse models, we evaluated the presence and clinical relevance of TMEM doorways in human PDAC (table1). Using immunohistochemical staining of FFPE sections from primary PDAC patient tumors, lymph nodes, and liver metastases, we identified TMEM doorways in all human PDAC specimens (Figure 2A-E and Table 1). Quantitative analysis demonstrated a significantly higher density of TMEM doorways in high-grade, poorly differentiated tumors (Figure 2C, H and I) compared with low grade, well-to-moderately differentiated (P < 0.05) (Figure 2D, H and I). Furthermore, tumors from patients who underwent up-front surgery for their PDAC had a median TMEM doorway density of 80 (65-105 count/10 high power fields) (Figure 2J and E). Tumors from patients who received FOLFIRINOX neoadjuvant therapy had a median TMEM doorway density of 30 (20–45) (P < 0.01) (Figure 2E and K).

**Figure 2.**
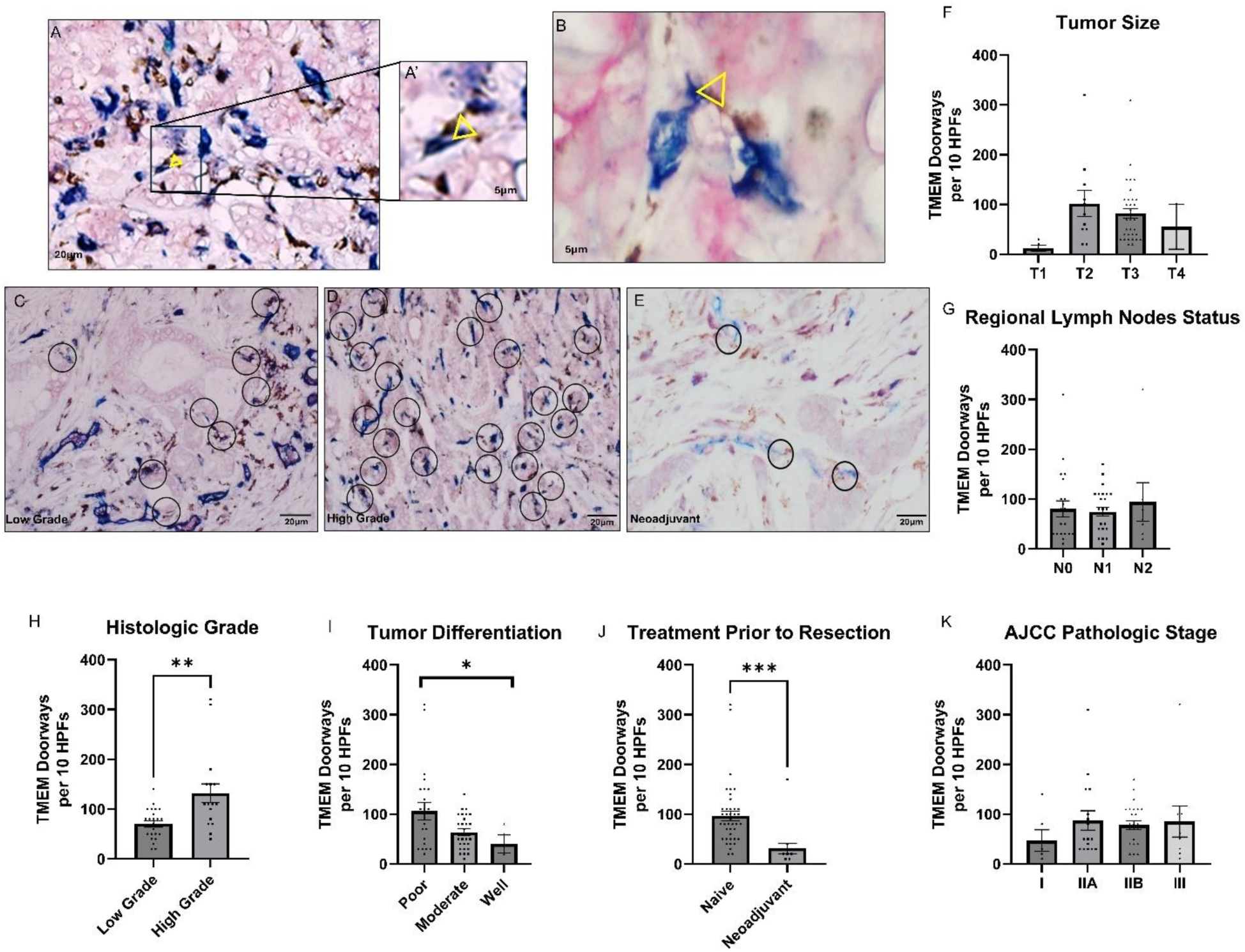
TMEM Doorways Are Abuntant in Human PDAC and Associated with Aggressive Tumor Features. (A-E) Triple immunostaining of TMEM doorways, macrophages are brown, endothelial cells are blue, and tumor cells expressing Mena are pink. (A and B) Representative image of TMEM doorway staining in primary PDAC tumor (A) and Liver met (B). In the enlarged image of the primary PDAC tumor (A’), and in the Liver met (B), the triangle denotes the three cells of the TMEM doorway, the macrophage is in direct contact with both the endothelial cell and the tumor cell. (C-E) Representative images of TMEM doorways (circles) in histologic sections: (C) Low grade PDAC tumors, (D) High grade PDAC tumors and (E) after Neoadjuvant therapy. Scale bar 20µm, circles 60 µm in diameter. (F-K) Distribution of TMEM doorway scores according to clinical pathological features in patients with PDAC: (F) Tumor size (T1: Tumor ≤ 2 cm, confined to pancreas. T2: Tumor > 2 cm and ≤ 4 cm. T3: Tumor > 4 cm, no major arterial involvement. T4: Tumor involving major arterial structure. (G) Regional Lymph Node Status(N0: No regional lymph node metastasis. N1: Metastasis to 1–3 regional lymph nodes. N2: Metastasis to ≥4 regional lymph nodes) H) Histologic Grade, (I) Tumor Differentiation, (K) AJCC Pathological Stage (Stage I: T1–T2, N0. Stage IIA: T3, N0. Stage IIB: Any T with N1. Stage III: Any T with N2 or T4 tumors involving major arteries) (J) Treatment prior resection. (* p < 0.05) analyzed by Mann–Whitney/Kruskal–Wallis.

**Table 1.** Clinicopathologic correlations of TMEM doorway scores in 60 PDAC patients.

| Variable | Subgroup | N (%) | Median TMEM (IQR) | p-value |
| --- | --- | --- | --- | --- |
| Gender | Male | 25 (41.7) | 80 (65–100) | 0.078 |
|  | Female | 35 (58.3) | 50 (35–75) |  |
| Race/Ethnicity | Non-Hispanic White | 8 (13.6) | 40 (30–50) | 0.779 |
|  | Non-Hispanic Black | 22 (37.3) | 80 (60–95) |  |
|  | Hispanic | 10 (16.9) | 50 (35–65) |  |
|  | Other | 19 (32.2) | 80 (55–105) |  |
| AJCC Stage | I | 8 (13.6) | 25 (20–40) | 0.63 |
|  | IIA | 17 (28.8) | 50 (35–70) |  |
|  | IIB | 25 (42.4) | 80 (60–100) |  |
|  | III | 9 (15.3) | 55 (40–80) |  |
| Tumor Size (T) | T1 (<2 cm) | 6 (10.2) | 15 (10–25) | 0.33 |
|  | T2 (2–4 cm) | 12 (20.3) | 70 (60–85) |  |
|  | T3 (>4 cm) | 39 (66.1) | 80 (65–105) |  |
|  | T4 (vascular involvement) | 2 (3.4) | 55 (50–60) |  |
| Lymph Node Status | N0 | 25 (42.4) | 50 (35–75) | 0.85 |
|  | N1 | 27 (45.8) | 75 (60–95) |  |
|  | N2 | 7 (11.9) | 60 (45–80) |  |
| Tumor Grade | Low | 24 (58.5) | 80 (65–100) | 0.008 |
|  | High | 17 (41.5) | 110 (90–130) |  |
| Tumor Differentiation | Poor | 28 (50.9) | 100 (80–115) | 0.05 |
|  | Moderate | 23 (41.8) | 50 (35–75) |  |
|  | Well | 4 (7.3) | 40 (35–45) |  |
| **Treatment** | **Naïve** | **41 (68.3)** | **80 (65–105)** | **<0.001** |
|  | **Neoadjuvant** | **19 (31.7)** | **30 (20–45)** |  |

\*Values represent median TMEM scores with interquartile ranges (IQR) for each subgroup.

In this cohort of patients who underwent surgery for their primary PDAC, tumor TMEM doorway density did not correlate with tumor size (Figure 2F), lymph node status (Figure 2G), or cancer stage (Figure 2J). The human cohort size was not large enough to stratify overall survival or metastasis-free survival based on TMEM doorway density.

### TMEM Doorways and Tie2-Expressing Macrophages in PDAC: Tie2 Inhibition and Intravasation

The validated IHC staining protocol for TMEM doorways includes staining for Mena, CD68, and CD31 (6, 10, 11). To further characterize the TMEM doorway associated macrophage and actionable signaling targets of the PDAC TMEM doorway, we multiplex stained murine PDAC tissues for IBA1, Tie2, and VEGF-A on sequential slides of TMEM doorway staining and overlaid the two staining sections. Analysis revealed that TMEM doorways predominantly contain Tie2-expressing macrophages (Figure 3A). In the KPCY T-low cell line model which has a robust macrophage tumor infiltrate, approximately 60% of identified TMEM doorways contained Tie2+ macrophages (Figure 3B). In the KPC model, Tie2+ macrophages were even more pronounced, present in over 90% of TMEM doorways (Figure 3C). Immunofluorescence analysis (Figures 3B and C) and flow cytometry (Supplemental Figure 1) revealed Tie2 expression on a variety of cell populations in PDAC (2% Tumor cells, 3% endothelial cells). Among tumor-infiltrating myeloid cells, Tie2 expression was confined to a small subpopulation (∼2%) of macrophages. Considering the immunofluorescent data showing that 60-90% of TMEM doorways have Tie2+ macrophages while the entire tumor mass has only 2% Tie2+ macrophages, there is a selective enrichment and spatial localization of Tie2+ macrophages at TMEM doorways within the TME.

**Figure 3.**
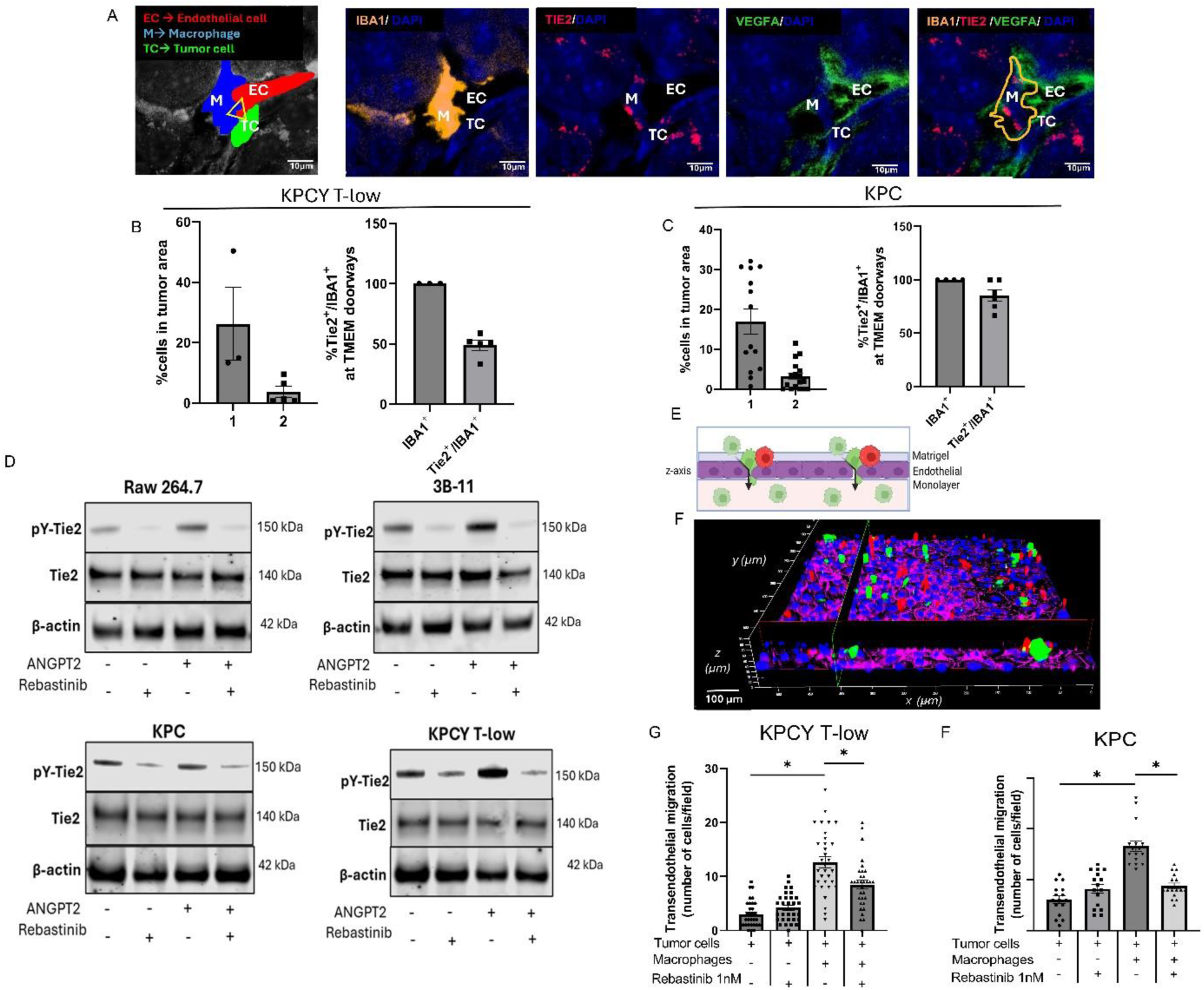
TMEM Doorways contain Tie2-Expressing Macrophages in PDAC: Tie2 Inhibition and Intravasation. (A) Immunofluorescence imaging of Tie2Hi/VEGFAHi macrophages in TMEM doorways in sequential sections. Endothelial cells (EC), macrophage (M) and tumor tissue (TC). In the merge panel, a mask for IBA1 macrophage was created in ImageJ^®^ for a better visualization of Tie2. Scale bar,10μm. (B and C) Quantification of Tie2+ macrophages in tumor area and withim TMEM doorway area of KPCY T-low (B) and KPC (C) tumors. (1)-total macrophages. (2)-Tie2 positive macrophages. (D) Western blot analysis of phospho-Tie2 immunoprecipitated from extracts of KPC, KPCY T-low, RAW 264.7 or 3B-11 cells treated with Rebastinib +/- Angiopoietin 2. (E) Experimental design of the subluminal to luminal trans-endothelial migration (iTEM) assay. Macrophages = red. Tumor cells = green. The tumor cells (green) are considered transendothelial migrating cells when they are crossing the endothelial layer (pink). (F) Representative image of iTEM assay from confocal microscope. Macrophages = red. Tumor cells = green. Endothelial layer = pink. Quantitation of trans-endothelial migration of KPCY T-low (G) and KPC (H). In KPCY T-low and KPC, we observed an increase of transendothelial migration from 3 cells/field to 12.65 cells/field and 6 cells/field to 16.65 cells/field respectively, in the presence of macrophages (p<0.05), following by a decrease to 8.46 cells/field and 8.46 cells/field after rebastinib treatment (p<0.05). Co-culture with macrophages significantly increased migration, which was inhibited by the Tie2 inhibitor rebastinib. Data are presented as mean ± SEM; (n ≥ 10 per group). *P < 0.05, by one-way ANOVA with Tukey’s post hoc test.

These findings led us to hypothesize that, as in breast cancer, Tie2 signaling in macrophages plays a functional role in mediating tumor cell intravasation (10, 12)(7, 28). To test this hypothesis, we examined the pharmacologic inhibition of Tie2 using rebastinib, a selective allosteric Tie2 inhibitor (12, 17). Immunoprecipitation of tyrosine phosphorylated Tie2 confirmed that rebastinib treatment (1 nM) effectively suppressed tyrosine phosphorylation levels of Tie2 in macrophages (RAW 264.7), endothelial cells (3B-11), as well as in PDAC cells (KPC and KPCY T-low) (Figure 3D). Dose-response assays confirmed that treatment with 1 nM rebastinib did not affect cell viability, ensuring that observed effects were not due to cytotoxicity (Supplemental Figure 2).

Because we found no cytotoxicity with rebastinib in tumor cells, endothelial cells, or macrophages and identified Tie2 expression in each of these cells *in vivo* and *in vitro*, we sought to identify Tie2’s functional role in PDAC intravasation (Figure 3E - H). In order to evaluate for transendothelial migration mechanisms in vitro, we sought to recapitulate TMEM doorway assembly and function using a stratified co-culture methodology (iTEM assay) (9, 18).

Transendothelial migration of PDAC cells was enhanced by macrophage co-culture. Notably, rebastinib treatment (1nM) significantly impaired macrophage-mediated tumor cell transendothelial migration (Figure 3E-H). Although Tie2 is expressed by tumor cells, macrophages, and endothelial cells, this functional assay showed *in vitro* transendothelial migration is mediated by macrophage Tie2 signaling.

### *In Vivo* Tie2 Inhibition Selectively Disrupts TMEM Doorway Activity and Tumor Cell Dissemination in PDAC

To investigate the functional role of Tie2⁺ macrophages *in vivo*, we evaluated the impact of pharmacologic Tie2 inhibition on TMEM doorway activity and metastatic dissemination using orthotopic PDAC models. Tumors, liver, lung, and peripheral blood were harvested for endpoint analyses. Immunohistochemistry was used to quantify TMEM doorways, while adjacent serial sections were analyzed by immunofluorescence to assess TMEM doorway activity, defined by the presence of extravascular HMWD adjacent to TMEM doorways, and Tie2^+^ macrophages (Figure 4A). The overall number of TMEM doorways remained unchanged following rebastinib treatment in both models (KPCY T-low and KPC) (Figure 4B and D). Importantly, a significant reduction in open (active) Tie2^+^ TMEM doorways was observed in KPCY T-low tumors (69.27% to 44.18%, p<0.05) and KPC tumors (90.88% to 69.12%, p<0.05) in Tie2+ TMEM doorways, as evidenced by decreased extravascular HMWD at Tie2^+^ TMEM doorways with rebastinib treatment.

**Figure 4.**
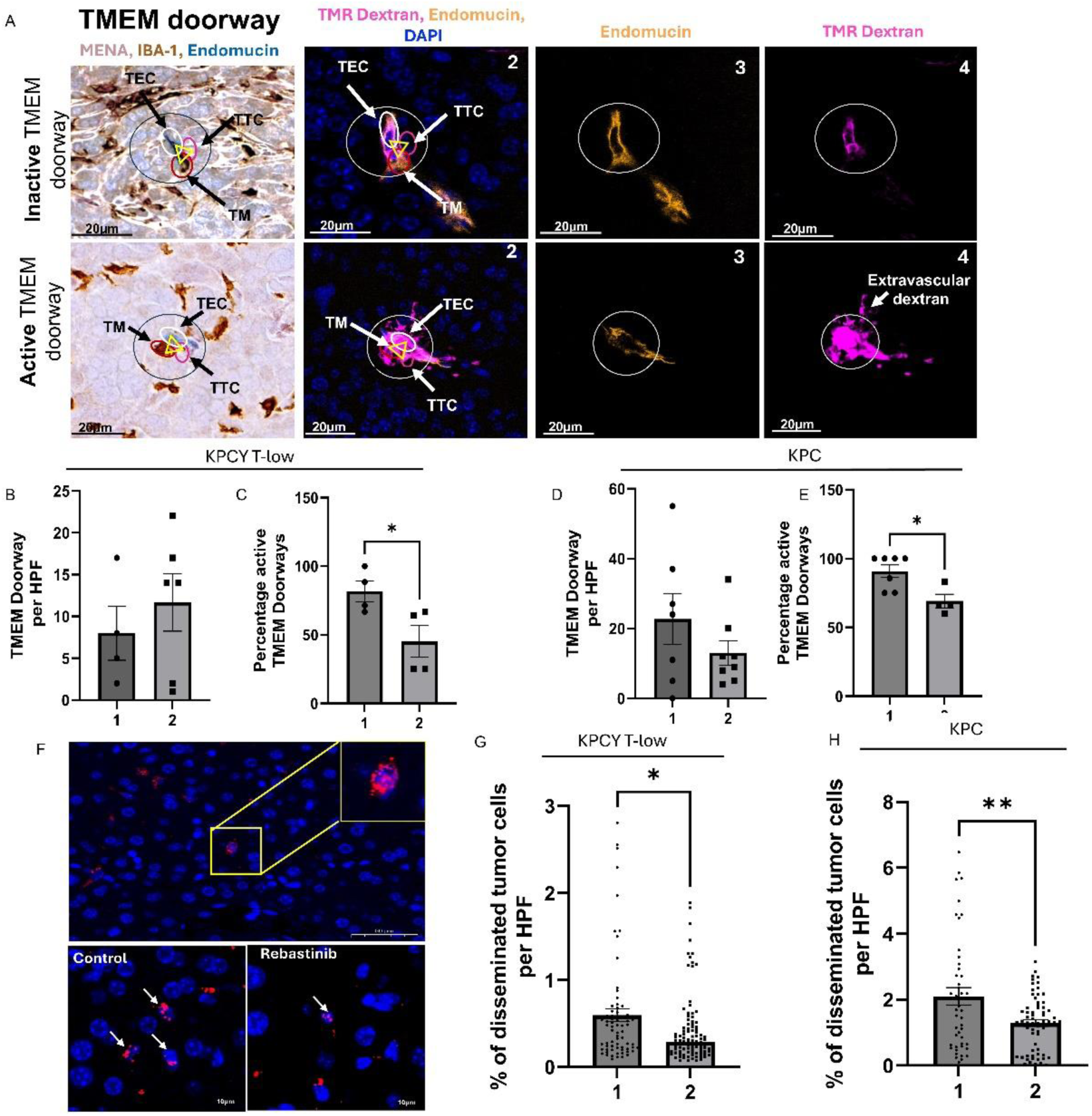
In Vivo Tie2 Inhibition Selectively Disrupts TMEM Doorway Activity and Tumor Cell Dissemination in PDAC. (A) panel 1 shows TMEM doorways, visualized by immunohistochemistry (IHC) staining for Mena, Iba-1, and endomucin. The three cells of the TMEM doorway (contained in black circle, with the three cells forming the TMEM doorway indicated with the yellow triangle) are the TMEM doorway endothelial cell (TEC, endomucin stained in blue), TMEM doorway macrophage (TM, Iba1 stained in brown), and TMEM doorway tumor cell (TTC, Mena stained in pink). Panels 2-4 show different staining channels of the sequential tissue section 2 aligned with the IHC stained section (panel 1) with immunofluorescence (IF) with antibodies against endomucin (panel 3 yellow), dextran (panel 4 pink) and nuclear stain DAPI (blue). Active versus inactive TMEM doorways were distinguished by the presence of extravascular dextran staining (panel), which indicates that the vessel had a TMEM doorway-associated vascular opening (TAVO). Scale bars=20μm. (B-H) Mice were treated with (1) control chow or (2) ∼0.44 mg/day of rebastinib for 3 weeks. (B and D) Quantification of TMEM doorway density in 10 high-power fields (HPFs; no significant difference) in KPCY T-low (B) and KPC tumors (D). (C and E) Immunofluorescence measurement of extravascular dextran levels in Tie2^+^ TMEM doorways indicate significant inhibition of TMEM doorways by rebastinib inKPCY T – low tumors (C) and KPC tumors (E). TMEM doorways were identified in the IF-stained sections as described above. Active TMEM doorways were identified by the presence of extravascular dextran which is not present in inactive TMEM doorways. *p<0.05. (F) Disseminated tumor cells in the liver stained with p53 (red). Increased magnification of field of the p53 positive cells is shown in the yellow box. The left bottom is a representative image of control and right bottom of rebastinib treated group in magnified fields. (G-H) Quantification of DTCs in the liver of animals as compared to the (1) control with (2) rebastinib treated groups showing significant inhibition of DTC levels. (G) KPCY T-low (H) KPC where each dot is a separate mouse. *p<0.05, analyzed by Student’s t-test. N=8.

Systemic dissemination was assessed by histological analysis of disseminated tumor cells (DTCs) in distant organs. Notably, rebastinib treatment reduced hepatic DTC burden in both models KPCY T-low (0.59% to 0.41% DTCs/field; p<0.05) and KPC (2.10% to 1.29% DTCs/field; p<0.05) underscoring its efficacy in reducing metastatic seeding (Figure 4F-H).

Importantly, quantitative analysis of the tumor microenvironment revealed no significant differences in either tumor-associated macrophage density (IBA1⁺ area) or microvascular density (endomucin⁺ area) between control and rebastinib-treated groups (Supplemental Figure 3). These findings suggest rebastinib’s mechanism of action *in vivo* is not anti-angiogenic nor related to vascular normalization (Supplemental Figure 3). Rather, these results indicate that Tie2 inhibition with rebastinib decreases TMEM doorway opening in PDAC.

### Perioperative Tie2 Inhibition in Combination with FOLFIRINOX enhances survival in an Orthotopic PDAC Model

To assess the therapeutic efficacy of Tie2 inhibition in PDAC, we conducted a survival study using an orthotopic KPCY T-low mouse model, incorporating surgical resection and chemotherapy to mimic a perioperative treatment strategy which would allow for evaluation of biologic end points as well as metastasis-related survival (Figure 5A).

**Figure 5.**
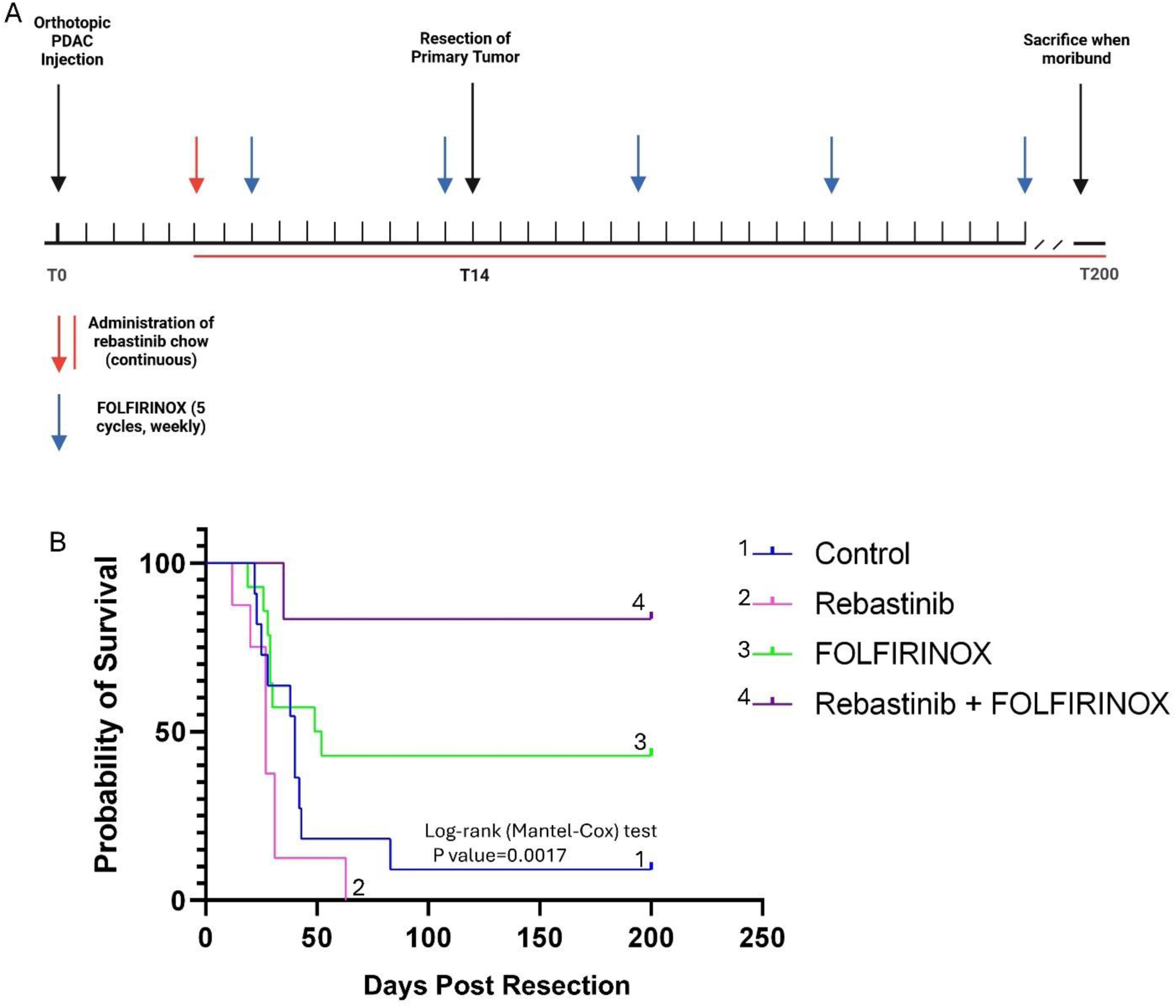
Tie2 Inhibition in Combination with FOLFIRINOX enhances survival in an Orthotopic PDAC mouse Model. (A) Schematic representing mouse treatment and monitoring. Red arrow indicates when rebastinib treatment started (day 5). The horizon red line represents a continuous treatment with rebastinib. Blue arrows indicate cycles of FOLFIRINOX treatment. T0 represents day 0, T14 – day 14 and T200 day 200. If the mouse was still live on day 200 it was sacrificed. (B) Intermittent treatment of KPCY T-low mice with rebastinib extends overall survival in combination with FOLFIRINOX, following the resection of the primary tumor. Primary PDAC tumors were resected on day 15 and treatment began on day 5. Dosing continued for 28 weeks and animals followed for survival. Control (n=10 mice), rebastinib (n=11), FOLFIRINOX (n=7), combo (n=6). P value is significant when comparing curves 1 and 4.

Primary tumors were evaluated pathologically (Supplemental Figure 4). All mice underwent complete gross and microscopic removal of their primary tumor (R0). All pancreatic transection margins were evaluated by a pathologist (PP) and negative for tumor cells.

No tumor had complete pathologic response. Partial pathologic response was evident in 4/8 rebastinib treated mice, 5/7 FOLFIRINOX treated mice, and 6/6 rebastinib and FOLFIRINOX treated mice (Supplemental table 1).

Necropsy of moribund mice revealed unique recurrence patterns in each treatment group. Nine of 10 control mice had recurrence. No recurrence was observed in 2/11 rebastinib treated mice, 3/7 FOLFIRINOX treated mice, and 5/6 rebastinib and FOLFIRINOX treated mice (Supplemental table 1). Patterns of recurrence revealed that most mice died of multisite (liver/peritoneum/regional) metastasis (5/10 in control mice, 4/11 in rebastinib treated mice) or regional lymph node metastasis. For FOLFIRINOX treated mice, recurrence was seen regionally in 3/7 with no liver metastasis (Supplemental table 1). For mice treated with combined FOLFIRINOX and rebastinib, one mouse had regional recurrence while no other mouse had metastatic recurrence (Supplemental Table 1).

Overall survival following curative intent resection did not significantly increase with rebastinib alone (overall survival 27 days - p = 0.133). However, FOLFIRINOX monotherapy yielded a modest, yet not statistically significant survival benefit (overall survival 50.5 days). The combination of FOLFIRINOX and rebastinib (overall survival not reached) resulted in a statistically significant extension in survival relative to rebastinib alone (overall survival 27 days) (p = 0.0082). Compared to untreated controls (overall survival 40 days), the combination of rebastinib and FOLFIRINOX showed a significant improvement in overall survival (p = 0.0017) (Figure 5B). This survival study was not powered to evaluate the differences in recurrence patterns but suggests that metastatic progression is decreased with combined perioperative FOLFIRINOX and rebastinib.

## Discussion

The diagnosis of PDAC has a poor prognosis and is rarely cured, due to its aggressive metastatic progression and resistance to therapies (1). While metastatic progression is often attributed to tumor-intrinsic properties, increasing evidence suggests a role for the tumor microenvironment in orchestrating the metastatic cascade, especially intravasation (2, 3) Our results show that intravasation of tumor cells in PDAC is an orchestrated mechanism that is executed by the TMEM doorway.

For the first time, we have observed the intravasation event in PDAC which appears to be dependent on a Tie2+ TAM. Migration and intravasation appear to be related to single cell migration and intravasation. Our results implicate the TMEM doorway as a mechanism for PDAC intravasation and dissemination. We also show improved survival and decreased metastatic burden in mice treated with combined Tie2 blockade resulting in inhibition of TMEM doorway opening, and cytotoxic chemotherapy.

The TMEM doorway is a portal of intravasation described in breast cancer (6, 12, 19, 20). At TMEM doorways, signaling between the associated cells causes the endothelial cell junction at the TMEM doorway to dissociate, allowing intravasation of tumor cells, and, as a result, hematogenous dissemination of tumor cells to secondary sites (20, 21) Initiation of TMEM doorway activity results from localized VEGF-A release from Tie2-expressing macrophages, which locally reduces cohesion of capillary endothelial adherents and tight junctions, thereby allowing a transient opening of the endothelial wall (∼30 min “opening”) (7). This opening is not related to vascular “leaking”. It is a highly localized, and transient, opening in the tumor endothelial layer of sufficient size to allow passage of an intact tumor cell, called TAVO (7, 9).

Our study provides the first evidence that TMEM doorways also exist and are operational in PDAC. Using high-resolution intravital microscopy in genetically engineered mouse models, we observed TAVO at perivascular macrophage-rich sites. These regions are similar to the TMEM doorways described in breast cancer, consisting of a MENA^High^ tumor cell, a Tie2-expressing macrophage, and an endothelial cell in direct contact (7, 10, 11). The visualization of tumor cell intravasation in real-time and 3D reconstruction of these tri-cellular structures confirm intravasation sites are TMEM doorways, thereby extending the relevance of this mechanism of intravasation to PDAC and supporting the hypothesis of a common mechanism of intravasation in carcinomas.

Importantly, our immunohistochemical analysis of human PDAC specimens revealed that TMEM doorways are present in all PDAC tumors including primary and metastatic sites. This implicates the TMEM doorway in initial metastatic dissemination but also in metastasis of metastases. Therefore, targeting this mechanism may reduce dissemination in primary and metastatic sites delaying the evolution of overwhelming metastasis. Since TMEM doorway density is variable in PDAC, TMEM doorways may serve as clinically relevant biomarker for metastatic risk and as an actionable target to disrupt dissemination in PDAC. These findings are aligned with prior reports in breast cancer where TMEM doorway density serves as an independent prognostic biomarker for distant metastasis in ER+/HER2− patients (6, 10, 11)

A defining feature of the TMEM doorway is the Tie2-expressing macrophage (7, 20, 21). Tie2 is expressed on endothelial cells and some epithelial cells and is primarily known for its involvement in vascular biology including angiogenesis (22). Our studies demonstrate that Tie2-expressing macrophages are enriched at TMEM doorways in PDAC. These cells, though representing a minority population (∼2%) of TAMs, are selectively enriched at TMEM doorways and display high VEGFA expression, consistent with their ability to modulate endothelial opening (7, 20, 23) (Figure 3A-C).

In breast cancer, the Tie2 blockade by rebastinib results in inhibition of tumor growth, invasion and metastasis. Examination of the effects of rebastinib at the cellular level demonstrates that rebastinib reduced tumor vascular density, and Tie2+ macrophages in the PyMT mammary tumor and its stroma (20, 21). In PDAC, we showed that pharmacologic inhibition of Tie2 signaling using rebastinib decreased Tie2 phosphorylation in both macrophages and endothelial cells and significantly impaired tumor cell transendothelial migration in co-culture assays (Figure 3D-H). In vivo, rebastinib selectively disrupted TMEM doorway activity without altering macrophage density or microvascular architecture (Supplementary Figure 2). This was associated with a reduction in disseminated tumor cells (DTCs) in liver tissue, further supporting the role of Tie2+ macrophages in intravasation (Figure 4). These results provide compelling evidence that Tie2-expressing macrophages are critical facilitators of TMEM doorway-mediated intravasation and metastatic dissemination in PDAC.

Although rebastinib monotherapy did not significantly prolong survival, its combination with the chemotherapy FOLFIRINOX resulted in a statistically significant survival benefit (Figure 5B). This combinatorial effect suggests that Tie2 inhibition may sensitize the metastatic niche to chemotherapy, likely by reducing the influx of new metastatic cells rather than directly affecting primary tumor growth. Perhaps inhibiting the TMEM doorway by Tie2 blockade blocks the educational niche for chemo-resistance and/or invasiveness, stemness, and dormancy and allows for sensitizing PDAC to FOLFIRNOX (24, 25). Taken together, these observations support the rationale for combining Tie2 inhibitors into treatment regiments for locoregional and metastatic stages of PDAC (26).

To illustrate our hypothesis in Figure 6, we proposed the mechanism by which TMEM doorways facilitate PDAC dissemination and how Tie2 blockade modulates this process. In panel 1, we show a perivascular Tie2+ macrophage engaging with a MENA^High^ tumor cell and an adjacent endothelial cell, forming the TMEM doorway structure. Panel 2 highlights the occurrence of a TAVO through which tumor cells intravasate, contributing to systemic dissemination. In panel 3, rebastinib-mediated inhibition of Tie2-signaling disrupts TMEM doorway function, reducing TAVO formation and thereby limiting intravasation, and distant metastasis. This model reinforces the conclusion that Tie2+ macrophages are a functional target in the metastatic cascade of PDAC.

**Figure 6.**
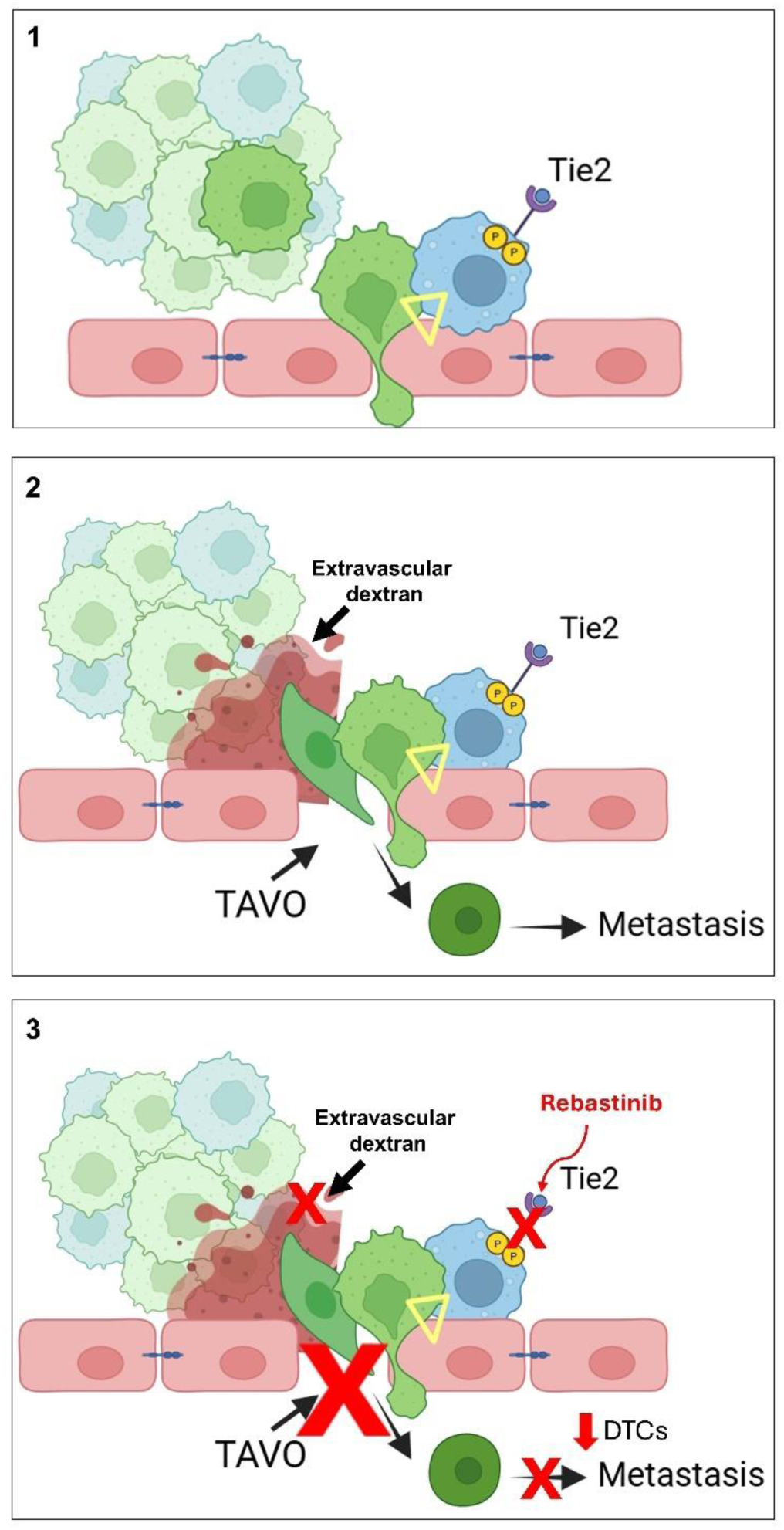
Inhibition of Tie-2 receptor with rebastinib can block TMEM doorway function and consequent intravasation, dissemination of tumor cells, and metastasis. Panel 1: The yellow triangle indicates a TMEM doorway containing a perivascular Tie2+ macrophage (blue) bound to a MENAHigh tumor cell (green), both tightly bound to an endothelial cell (pink). Panel 2: shows the TMEM doorway is associated with transient opening of the endothelial layer resulting in TMEM doorway associated vascular opening (TAVO) through which tumor cells (green) intravasate, contributing to systemic dissemination. Panel 3: rebastinib-mediated inhibition of Tie2 signaling inhibits TMEM doorway function, reducing TAVO (red x) thereby limiting intravasation, and distant metastasis.

Collectively, our study identifies TMEM doorways as operational TME structures in PDAC and establishes Tie2+ macrophages as facilitators of tumor cell intravasation. The efficacy of rebastinib in combination with FOLFIRINOX provides strong preclinical evidence for clinical trials evaluating Tie2 inhibition untapped therapeutic opportunity in the PDAC space. Whether the TMEM doorway density serves as a predictive biomarker for patient stratification and therapeutic response as demonstrated in ER+/HER2− breast cancer is unknown and requires a larger patient cohort to evaluate (6, 10, 11).

The molecular mechanisms of TMEM doorway assembly in PDAC and the role of stromal heterogeneity in modulating their activity is still unclear and requires further study. We have shown that Tie2 inhibition disrupts dissemination and chemotherapy efficacy, establishing TMEM doorways as potential therapeutic targets and predictive biomarkers of metastatic risk in PDAC.

## Materials and Methods

### Mice

All mouse studies were conducted in accordance with the National Institutes of Health regulations concerning the care and use of experimental animals and approved by Albert Einstein College of Medicine Institute for Animal Care and Use Committee (Protocols: 00001845 and 00001079). The C57/BL6 (000664 – Jackson Laboratory) strain was used for pancreatic cancer orthotopic injections and tissue collection. Each treatment group size has a minimum of 8 mice.

For intravital imaging we used three models: (1) C57B6: Csf1r-GAL4-VP16/UAS-ECFP (ECFP labelled macrophages) (JAX:026051) orthotopically injected with KPCY T-low cells (27); (2) KrasG12D/p53fl/+/Pdx1-Cre/RosaYFP (KPCY, YFP labelled pancreata that spontaneously forms pancreatic tumors); (3) Rag2KO (JAX:008449) Csf1r-GAL4-VP16/UAS-ECFP (ECFP labelled macrophages) orthotopically injected with C-EMT 3077 cells (28–30). Mice were housed in static cages under specific pathogen-free conditions in a temperature- and humidity-controlled environment. All mice were provided with 1/8-inch corn cob as bedding material. All experiments were performed during the daytime of a 12-hour day/night cycle. KPCY mice were used between 16 and 20 weeks of age, when they spontaneously develop tumors.

To generate the orthotopic tumors we used two different cell lines, the KPC cells and KPCY T-low cells, a total of 6000 cells per animal were re-suspended in 50ul of sterile PBS and injected in the tail of the pancreas under general anesthesia by way of a left subcostal abdominal incision during survival surgery (31). Primary tumors developed over 21 days (KPCY T-low) or 28 days (KPC). After 5 days of injection of tumor cells, mice were fed with either control chow or chow supplemented with rebastinib (DCC3014-110 mg/kg, Deciphera™) with a calculated 0.44mg/day rebastinib dose as previously described (32).

All mice underwent a R0 distal pancreatectomy to completely remove the primary tumor on post tumor cell injection day 15 and subsequently monitored. For our overall survival study, 5 days after orthotopic tumor implantation, mice were randomized into four treatment arms: control, FOLFIRINOX alone, rebastinib alone, or a combination of rebastinib and FOLFIRINOX. Rebastinib, was administered continuously via chow beginning on day 5. FOLFIRINOX was delivered as two cycles pre-resection and three cycles post-resection. The dose of each drug was weight based – 5-FU (33mg/kg), Irinotecan (33 mg/kg) and Leucovorin (67mg/kg) OXALIPLATIN (3mg/kg) (33).

### Intravital Imaging

All imaging was performed on a previously described, custom-built, inverted two-laser multiphoton microscope (23) or using an Olympus FluoView 1000 system equipped with an inverted IX81 microscope body. A Mai Tai laser was tuned to 960 nm. Imaging was conducted with a 25× objective lens (numerical aperture 1.05). Emission signals were collected through the following bandpass filters: 450/70 nm (415–485 nm) for the blue channel, 520/35 nm (502.5– 537.5 nm) for the green channel, and 582/64 nm (550–614 nm) for the red channel. CFP fluorescence was detected in both the blue and green channels, while YFP was captured in the green channel. TRITC-dextran (155 kDa -cat# T1287, Sigma-Aldrich, Burlington, MA, USA), used as a vascular dye, was detected in the red channel. Images were acquired at 12-bit depth and 2 frames per second. The pancreatic window assessment and imaging was performed as described before (13, 14). The vasculature of the tissue was identified by circulating erythrocytes and serum labeled with TMR dextran. Tumor tissue was identified by YFP labeling. A minimal amount of image processing was performed to optimize brightness and contrast of images.

### Human specimens

PDAC samples were collected from patients who underwent curative-intent surgical resection between 2013 and 2022 at a single tertiary care institution (n=60, IRB-2018-8906). Patients were selected based on the presence or absence of subsequent metastases. For each patient, the diagnosis of PDAC was confirmed.

### Cells

KPC ( Cat.#: 153600: Cancer tools.org) (28), KPCY T-low (6419c5) (34), or KPCY C-EMT 3077 (provided by Ben Z. Stanger) (35), murine macrophage cell line RAW264.7 (ATCC-provided by Dianne Cox), and murine endothelial cells 3B11 (ATCC - provided by Camille Duran) were cultured in Dulbecco’s Modified Eagle Medium (DMEM) (cat# SH30243, Hyclone, GE Healthcare Life Sciences, Logan, UT, USA) supplemented with 10% fetal bovine serum (FBS) (cat# S11550, Atlanta Biologicals, Flowery Branch, GA, USA).

### Histology (H&E) and immunohistochemistry for TMEM doorways

After mice were sacrificed, all PDAC tumors were extracted and immersed in 10% formalin in a volume ratio of tumor to formalin of 1:7. Tissues were fixed for 24 to 48 h and embedded in paraffin, then processed for histological examination. One 5 µm section from each tumor was stained for hematoxylin and eosin (H&E) and one for TMEM doorways. TMEM doorway stain was performed as previously described (6), except that in this study we used anti-panMena antibody (510693; BD Biosciences) to detect Mena-expressing cancer cells. To visualize macrophages, we used anti-IBA1 antibody (019-19741; Wako) for mouse and anti-CD68 (MO876; Dako) for human tumors. To visualize endothelial cells, we used anti-endomucin (SC-65495; Santa Cruz) for mouse and anti-CD31 (MO823; Dako) for human tumors. TMEM doorways were counted in 10 high power (40x) fields, and the sum was used as the score called TMEM doorway density.

### Immunofluorescence labeling of tumor vasculature and extravasation with 155 kDa dextran-TMR

Labeling flowing vasculature and sites of permeability was performed as previously described (7). Briefly, to quantify extravasation, high molecular weight 155 kDa TMR-dextran (cat# T1287, Sigma-Aldrich, Burlington, MA, USA) diluted in PBS was administered via retro-orbital injection 15 minutes before the experiments to label patent blood vessels. Resected tumors were fixed for 48 hours in 10% formalin in a volume ratio of tumor to formalin of 1:7 and made into paraffin blocks. Paraffin blocks of tumors were cut into 5 μm sections and stained by using rabbit anti-TMR (A-6397; Life 169 Technologies, Carlsbad, CA, USA).

### Immunoprecipitation followed by western blotting

RAW 264.7 macrophages, 3B-11 endothelial cells, and KPC/KPCY pancreatic cancer cells were serum-starved (DMEM + 0.5% FBS), treated overnight with 1 nM Rebastinib or DMSO (1:10,000), and stimulated with 800 ng/mL Angiopoietin-2 with or without 1 NM Rebastinib for 15 minutes. Cells were lysed (Invitrogen Lysis Buffer II), and lysates (1 μg/μL in DPBS) were precleared with Protein G Sepharose beads, then incubated with 5 μg anti-phosphotyrosine antibody (4G10; Cell Signaling) or isotype control (2 h, 4°C). Immune complexes were captured with Protein G beads, pelleted (14,000 rpm, 4°C), resolved by SDS-PAGE (NuPAGE, Invitrogen), and immunoblotted for Tie2 (LSBio).

### Transendothelial Migration Assay (iTEM assay)

The iTEM assay was performed as previously described with modifications (8, 36) and briefly described here. The transwell (8 μm pore size; cat# 353097, Corning, Corning, NY, USA) was prepared so that tumor cell transendothelial migration was in the intravasation direction found in vivo (from subluminal side to luminal side of the endothelium). To prepare the endothelial monolayer, the underside of each transwell was coated with 50 μL of Matrigel (2.5 μg/mL; cat# 354230, Corning). Approximately 8×10^4^ 3B11 cells were plated on the Matrigel coated underside of the transwell. Transwells were then flipped into a 24-well plate containing 200 μL of DMEM and monolayers were formed over a 16 hours period. The integrity of the endothelium, defined as a confluent endothelial barrier, was measured using low molecular weight dextran as described previously Macrophages and tumor cells were labelled with cell tracker dyes (cat# C7025, C34552, Invitrogen) before each experiment. Then, 1.5×10^4^ tumor cells and 5×10^4^ macrophages were added to the upper chamber of the transwell in 200 μL of DMEM without serum while the bottom chamber contained DMEM supplemented with 36 μg/mL of CSF-1. At the time of seeding cells were treated with vehicle (DMSO) or rebastinib (1nM – DCC3014-DP-001919.TO.16- Deciphera Pharmaceuticals Inc, USA). After 18 hours, the transwells were fixed and stained for ZO-1 (cat# 402200, Invitrogen) as previously described. Transwells were imaged using a Leica SP8 confocal microscope using a 20× 1.4 NA objective and processed using Leica software. Tumor cell transendothelial migration quantification was performed by counting the number of tumor cells that were crossing the intact endothelium (intact monolayers confirmed by ZO-1 staining for tight junctions) within the same field of view (20x, 8 random fields) and represented as average values from at least three independent experiments.

### Immunofluorescence staining of tissue

For mouse specimens, tumor sections were dewaxed in xylene and rehydrated in alcohol followed by water. Antigen retrieval was performed with a citrate solution and blocked with a blocking solution (3% BSA in PBS). Serial sections from each tumor were stained for H&E and TMEM doorway. The sections were stained for: anti-endomucin (cat# sc-65495, Santa Cruz Biotechnology), anti-TMR (to visualize TMR-dextran, cat# 183 A-6397, Thermo Fisher Scientific), anti-Iba1 (cat# ab283346, Abcam), anti-Tie-2 (cat# LS-B14782-50, LSBio) and Anti-VEGF-A (cat# A121588, antibodies.com). The primary antibodies were detected with Alexa Fluor 488, 555 and 647 conjugated secondary antibodies (Invitrogen, Eugene, OR, USA), or Alexa 759 (Abcam, USA) and nuclei were stained with 4, 6-196 diamidino-2-phenylindole (DAPI). All fluorescently labeled samples were mounted with Prolong Diamond antifade reagent (cat# P36961, Invitrogen) and imaged with a PerkinElmer Pannoramic 250 Flash II digital whole-slide scanner using a 20x 0.8NA Plan-Apochromat objective (PerkinElmer, Hopkinton, MA, USA). Whole-tissue images were uploaded in Slide Viewer version 2.2 (3DHISTECH).

For human specimens, each block was serially sectioned and stained to identify TMEM doorways, following previously described protocols (6, 8, 10). Macrophages, tumors cells, and endothelial cells were identified as CD68+ (anti-CD68-mouse, Dako #MO876; DAB chromogen, Dako #K3468), pan-MENA+ (anti-panMena-mouse A351F7D9, MilliporeSigmwa #MAB2635; purple chromogen, Vector #SK-4605), and CD31+ (anti-CD31-rabbit, Abcam #ab182981; blue chromogen, Vector #AK-5001), respectively. The stained slides were imaged using the PerkinElmer Pannoramic 250 Flash II digital whole-slide scanner using a 20x 0.8NA Plan-Apochromat objective (PerkinElmer, Hopkinton, MA, USA). TMEM doorways in PDAC samples were scored by three pathologists (MO, NP, PP) according to established criteria (6, 10). TMEM doorway scoring was performed within tumor regions of interest, with each TMEM doorway marked by a 50μm-diameter circle in Adobe Photoshop ®. TMEM doorway scores were calculated as the total count within the tumor region of interest for each patient.

### Extravascular dextran and Tie-2 quantification at TMEM doorways

To identify active TMEM doorways, we utilized an automated method of a prior TMEM doorway activity assay that takes into account the presence of extravascular high-molecular weight dextran at TMEM doorways (TMEM doorway associated vascular opening, TAVO) (7, 8, 20).

In short, serial tissue sections were cut and stained for TMR-Dextran and endomucin by IF (tissue section 1) or IBA1, Tie-2, VEGF-A (tissue section 3) and for TMEM doorways by staining for Iba1, endomucin, and Mena by IHC (tissue section 2). The sections were then imaged with a digital whole slide scanner and aligned to the single cell level using the TissueAlign module in Vis (Visiopharm, Hoersholm, Denmark). Images were thresholded and masked for endomucin (blood vessel) and dextran signals. These masks were then superimposed to be able to differentiate between intravascular and extravascular dextran signal. In the IHC section, TMEM doorways were identified using previously published criteria (6, 10). In this manner, the extravascular dextran around TMEM doorways could be quantified. For quantification of IBA1^+^/Tie2^+^ macrophages the TMEM doorway associated tumor cell was identified using the IHC section, within the TMEM doorway circle, and the TMEM doorway spot was marked at the tissue section 3. Using the cell quant from Slide Viewer, a mask was created to quantify the IBA1^+^/Tie2^+^ inside of TMEM doorway, allowing the quantification of Tie2^+^ macrophages at TMEM doorways.

### Disseminated tumor cells (DTCs) in livers

To detect single disseminated tumor cells immunofluorescence staining for p53 clone POE316A/E9 (MABE1808, Millipore) was performed. The analysis was conducted using CellQuant from Slideviewer v2.2. For each case, 10 fields of view (mm) were selected and DTCs were counted based on nuclear detection of p53. The density of DTCs in the metastatic site was expressed as percentage of DTCs per total area of the liver tissue analyzed.

### Statistical analysis

Individual animals in each cohort are presented as individual points in a dot plot. A horizontal line indicates the mean value, and the error bars represent the standard deviation or standard error of the mean, as indicated in the figure legend. Statistical significance was determined using an unpaired Student’s t-test, one-way ANOVA, or two-way ANOVA, with Tukey’s multiple comparisons test, as indicated, using GraphPad Prism (version 10; Graph Pad 306 Software, La Jolla, CA). Correlation was determined using Pearson’s Correlation Coefficient. Data sets were checked for normality (D’Agostino & Pearson omnibus normality test or Shapiro-Wilk normality test) and unequal variance using GraphPad Prism. Welch’s correction was applied to t-tests as needed. P values less than 0.05 were deemed significant. For in vitro experiments, results are representative of at least three independent experiments. For human analysis, binary subgroup comparisons were performed using the Mann–Whitney U test; comparisons involving >2 subgroups used the Kruskal–Wallis H test (reported with degrees of freedom). Raw and FDR-adjusted p-values (Benjamini–Hochberg) are shown; statistically significant associations (FDR-adjusted p < 0.05) are highlighted in bold.

## Acknowledgments

This work was supported by the DOD PA210223P1-W81XWH, NIH R01-CA-276512 and NCI P30CA013330 grants, the shared instrumental grants S10OD026852-01A1 and 1S10OD023591-01, the Gruss-Lipper Biophotonics Center, the Integrated Imaging Program, the Integrated Imaging Program for Cancer Research, The Evelyn Gruss-Lipper Charitable Foundation. We would like to thank the Translational Pathology Laboratory for autostaining support. Hillary Guzik, Rotem Alon, Vera Desmarais and the Analytical Imaging Facility at Albert Einstein College of Medicine for imaging support. Suryansh Shukla for computer analyses supported. Histology and Comparative core for tissue processing. Dr. George Karagiannis and Dra. Dimitra Anastasiadou for technical insights and lab space. Dr. Julio Aguirre-Ghiso for editing and reviewing the manuscript.

## Author contributions

### Conceptualization

JCM, EPZ, DC, NCP, MHO, JSC. Methodology: JCM, EPZ, LAA, FPM, YQZ, JL, YJ, CA, JP, RE, PP, NCP, JSC. Formal Analysis: JCM, EPZ, LAA, XY, PP, NCP, DE, MHO, JSC Software: YJ, DE Investigation: EPZ, LAA, HGH, DC, JCM, BZS. Resources: JCM, BZS, MHO, JSC. Writing: EPZ, JCM, JSC. Funding Acquisition: JCM, BZS, MHO, JSC Supervision: JCM, BZS, JSC. All authors read and approved the submitted version and have agreed to be personally accountable for their own contributions and to ensure that questions related to the accuracy or integrity of any part of the work are appropriately investigated and resolved.

### Other interests

BZS has received grants from Boehringer-Ingelheim and Revolution Medicines and holds equity in iTeos Therapeutics.

