## Supplemental Figures for "TMEM Doorway Mediated Metastasis in Pancreatic Ductal Adenocarcinoma by Tie2 Signaling"

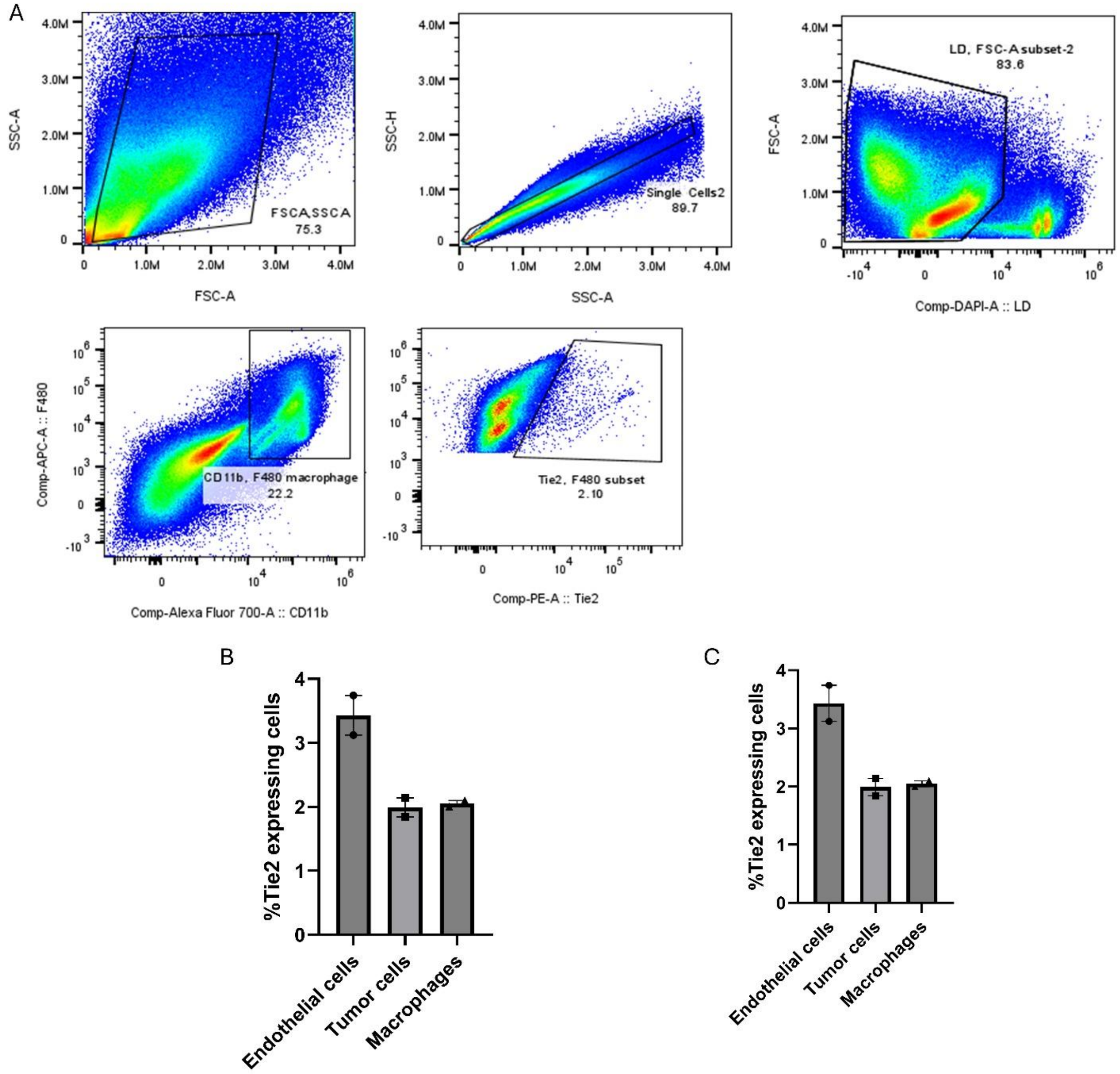


**Supplemental Figure 1. Flow cytometry showing Tie2 expression on a variety of cell population in PDAC**. (A) Gating strategy use to determine Tie2 in macrophages. (B) Tie2 expression in endothelial cells, tumor cells and macrophages. (C) Percentage of macrophages compared to other cell types in the tumor and the percentage of Tie2 expressing macrophages compared to other cell types in the tumor. Tie2 expressing macrophages are 2% of all macropages in the tumor.

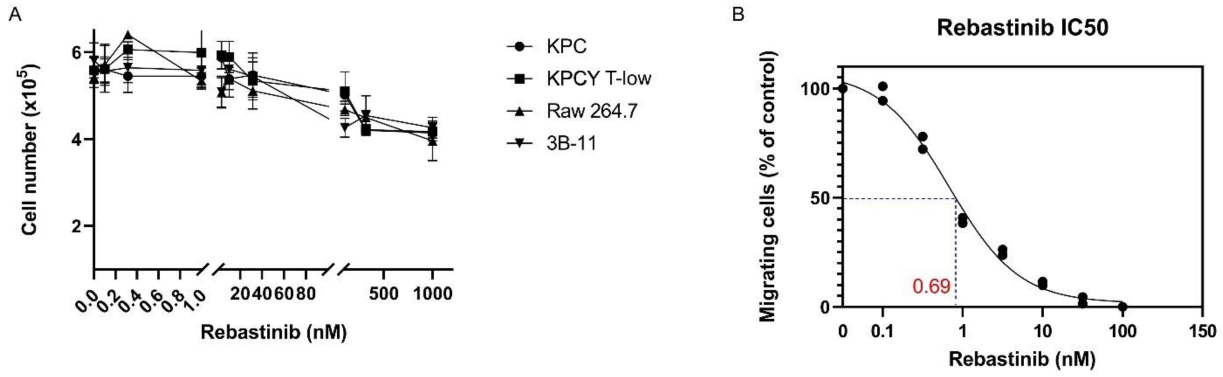


**Supplemental Figure 2**. **Rebastinib 1nM is not cytotoxic for cells.** (A) Toxicity assay of KPC, KPCY T-low, 3B11 and Raw 264.7 cells showing that at the concentration of 1nM Rebastinib is nontoxictoxic for all. (B) Graph showing that a concentration of 0.69nM of rebastinib in a iTEM assay has an inhibitory effect reducing 50% of transendothelial migrating cells. Data was analyzed using GraphPad Prism. N=3 replicates)

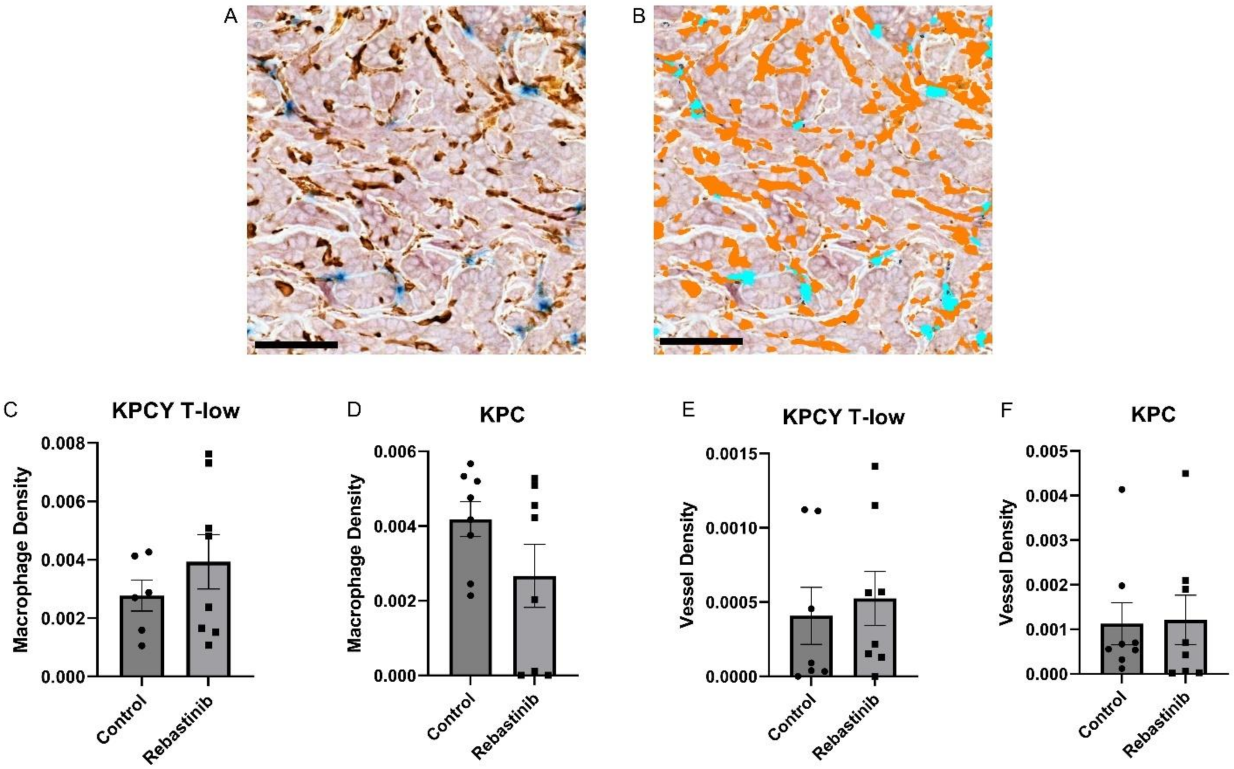


**Supplemental Figure 3**. **Rebastinib does not change macrophage infiltration or vessel density in pancreatic ductal adenocarcinoma.** (A-B) Representative images of macrophage and microvascular density quantification in PDAC TMEM doorway IHC stained. (A) Unprocessed TMEM doorway-stained images. Blue = Vessels (endomucin), Pink/Red = Tumor cells (panMena), Brown = Macrophages (IBA1). (B) Classified image with an overlay identifying pixels belonging to macrophages (brown) and vessels (blue). Scale bar = 50 µm (C-D) Quantification of macrophages shown in (C) KPCY T-low and (D) KPC tumors treated with control and rebastinib. (E-F) Quantification of vessel density shown in (E) KPCY T-low and (F) KPC tumors treated with control and rebastinib.

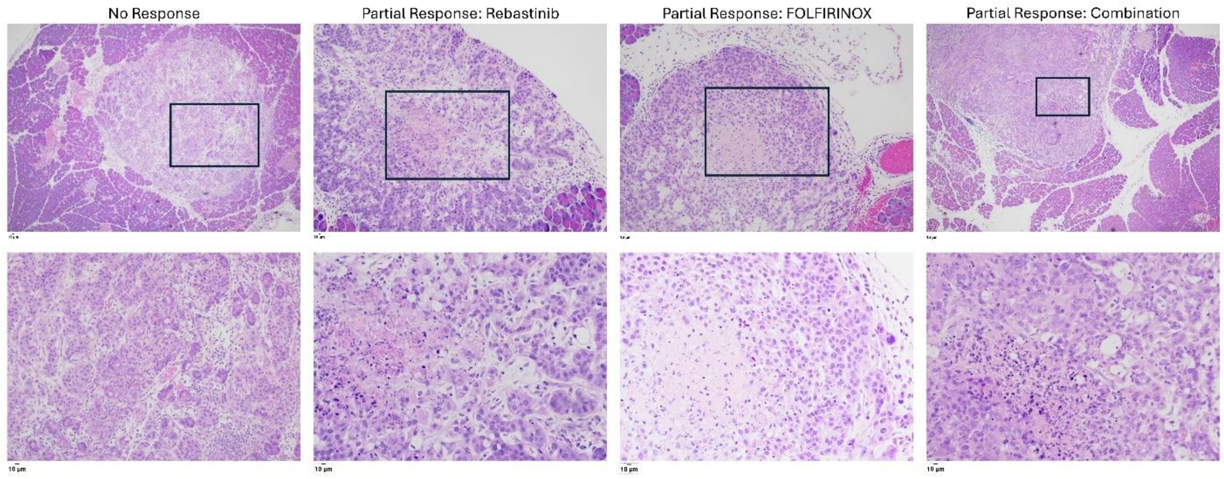


**Supplemental Figure 4. Resected tumors from mice of the survival study stained with hematoxilin/eosin.** First panel showing no response to treatment. Second panel showing partial response to Rebastinib. Third panel showing partial response to FOLFIRINOX. Fourth panel showing partial response from the combination (Rebastinib + FOLFIRINOX). Top panel 10X zoom. Bottom panels zoom on the black marked area in the top panels. The response was determined by the pathologist according to the College of American Pathologists (CAP) guidelines: CAP Grade 0 — Complete Response No viable residual tumor identified. Only fibrosis, mucin, or inflammation remains. CAP Grade 1 — Near Complete Response. Minimal residual cancer, with single cells or rare small groups of tumors. Robust therapy-induced fibrosis is present. CAP Grade 2 — Partial Response. Residual cancer is present, but there is evidence of tumor regression, such as fibrosis replacing parts of the tumor. CAP Grade 3 — Poor or No Response. Extensive residual cancer with minimal or no evidence of tumor regression. The results are described in supplemental table 1.
